# Thermodynamic Modeling of mRNA with the Addition of Precipitants

**DOI:** 10.1101/2025.10.18.683252

**Authors:** Vico Tenberg, Pierre J. Walker, Pavan K. Inguva, Maria del Carme Pons Royo, Andrew L. Acevedo, Vernon S. Lin, Marion Pang, Zhen-Gang Wang, Allan S. Myerson, Richard D. Braatz

## Abstract

Nucleic acid therapeutics (NATs) have recently emerged as an exciting therapeutic modality for a range of indications, most notably as vaccines for SARS-CoV-2. In many cases, the thermodynamics of a system containing nucleic acids (such as in the downstream purification of mRNA from solution or within the lipid nanoparticle) can significantly influence the properties and efficacy of that system. Consequently, an accurate thermodynamic description of the system is essential for understanding and optimizing that system. In this work, the SAFT-*γ* Mie equation of state was used to predictively model mRNA solubility. Experimental measurements of the solubility of two different mRNA sequences in various conditions (namely choice of precipitant(s), precipitant concentration, and temperature) were obtained and used to validate the model. Not only was the thermodynamic model able to quantitatively predict the solubility of mRNA in solution under different conditions, it was also able to yield mechanistic insight into the factor driving precipitation, namely the disruption of water-mRNA hydrogen bonding. The developed model can be extended to other mRNA sequences in a range of conditions beyond the experimental data presented in this work.

## Introduction

Nucleic acid therapeutics (NATs) are gaining immense interest as a drug modality for treating a wide range of indications. To that end, there is significant research and development (R&D) throughout the value chain to support their development and commercialization. One of the key enabling technologies driving R&D initiatives for NATs is the use of digital workflows and tools in various aspects ranging from sequence design /mRNA structure optimization [1, 2], lipid nanoparticle (LNP) formulation [3, 4], to NAT drug substance/product manufacturing [5, 6].

A key input to many of these tools, a rigorous description of the thermodynamics of nucleic acids and their interaction with other species, is lacking. This gap is not surprising, as mRNA is not only a large biological molecule, it contains the full spectrum of chemical interactions (dispersion, polarity, hydrogen bonding, and electrostatics) each of which is challenging to model individually, and even more so when they act simultaneously. Molecular modeling of mixtures containing nucleic acids would require computational methods with chemical specificity across scales, ranging from *ab initio* quantum calculations to molecular dynamics simulations [7–12]. While powerful for probing microscopic interactions, these methods face severe limitations in addressing macroscopic phenomena such as precipitation or in computing properties such as solubilities, due to computational cost and scalability. Data-driven approaches could in principle address such computational barriers. However, the chemical complexity of mRNA-containing systems, combined with the scarcity of reliable experimental solubility data, makes these approaches impractical at present. The utility of such data-driven models to predict the properties of nucleic acid mixtures in conditions outside the training dataset (i.e., to extrapolate) could also pose a significant challenge.

Within the context of NAT drug substance/product manufacturing specifically, we highlight three cases where a rigorous thermodynamic model of nucleic acids can be of immense value: 1) in vitro transcription (IVT) for RNA synthesis. In the IVT reaction, the kinetics of many reaction steps are a function of the equilibrium properties of magnesium (an enzyme cofactor), nucleoside triphosphates (feedstock), and pyrophosphate (a byproduct) [13, 14]. Describing the reaction kinetics at higher concentration regimes, relevant for process intensification and optimization, requires thermodynamic models that can accurately capture nonideal solution properties. 2) RNA separation and purification. In view of the inefficiency of chromatographic separation methods for large-scale purification, precipitation-based methods have emerged as an alternative separation method for purification of RNA from the IVT reaction mixture [15–17]. Detailed knowledge of the phase behavior of RNA in various process conditions (e.g., solvent composition, precipitant concentration, and temperature) is needed for process design and optimization. 3) LNP formation. The formation of LNPs can be described as a phase transition process where the lipids (dissolved in ethanol) and RNA (dissolved in an aqueous buffer) precipitate together upon mixing. Not only are the thermodynamics of the system critical in influencing LNP structure [18, 19], they are also a key input for various mechanistic models of LNP formation [6, 20]. We also anticipate such a model would be of use in other areas of biology and biotechnology such as better understanding biomolecular condensates (e.g., [21]).

To address this gap, we present a systematic experimental dataset of mRNA solubilities measured in various conditions (temperatures, precipitant concentrations, and mixed solvents) and a rigorous and predictive thermodynamic model for RNA that is validated against the dataset. For the thermodynamic model, the physics-based Statistical Associating Fluid Theory (SAFT) method, specifically the SAFT-*γ* Mie variant, is used. SAFT (and its variants) enables a rigorous statistical-mechanics-based description of the various interactions present and has been used successfully to describe complex systems such as polyelectrolytes [22] and amino acids [23]. In addition, some variants of SAFT, including SAFT-*γ* Mie, use a group-contribution approach to parameterization which enables their extension to a broad range of compounds and systems. To apply the SAFT-*γ* Mie model to nucleic acid systems, a significant number of new interaction parameters had to be fitted for the functional groups present (that were not available in previous implementations of the model), which required a careful parameterization strategy to ensure accuracy and transferability of the model.

## Materials and Methods

### Experimental Measurements

Two mRNA sequences are considered within this work, referred herein as FLuc (2117 bp, poly-30/70(A) tail, *A*_260*/*280_ = 2.00) and COVID (4283 bp, poly-30/70(A) tail, *A*_260*/*280_ = 1.74). The sequences examined in this work contain approximately equal distributions of the four nucleobases, with only short domains of identical bases. We note that N1-methyl-pseudouridine is used instead of uridine within both sequences. The sequences were delivered in aqueous nuclease-free solution with a concentration of 0.323 and 1.106 mg mL^*−*1^ for FLuc and COVID, respectively. These solutions were kept at *−*20° and *−*80°C for short- and long-term storage, respectively. To verify that no degradation occurred, these concentrations were validated via NanoDrop measurements prior to each experimental run. All experiments were performed using RNase-free vials and pipette tips to avoid contamination and/or degradation of the mRNA. Additionally, prior to every experimental run, bench surfaces, equipment, and personal gloves were cleaned with *RNase Away* ™, supplied by Thermo Fischer Scientific. Salt solutions of 20 wt%, for sodium acetate and ammonium acetate, and 7 wt% for sodium chloride, were prepared with RNase-free water. Similarly, aqueous PEG6k solutions were prepared with a concentration of 30 wt%.

To determine the solubilities of various mRNA/precipitant systems, 100–200 µL of mRNA solution was mixed with the desired amounts of salt solutions (shown visually in Figure 1a, left illustration) and additional precipitants such as ethanol or aqueous PEG6k solution. The vials were weighted with a *TB-215D* balance from Denver Instrument (*d* = 0.01 mg) after every subsequent addition of the components. Their weight was recorded to obtain the total composition of the mixture. The samples were left to equilibrate for 24 h. In preliminary experiments, it was shown that a minimum equilibration time of *∼*6 h is required and that the mRNA is stable in these conditions for the equilibration period of at least 24 h. This is demonstrated in Figure 1b where the solubility of FLuc in an aqueous mixture with added sodium chloride reaches its final value around *∼*6 h and does not change significantly even after additional 20 h. Subsequently, the solutions were stirred at 20 rpm on a revolving rotator in a cold room at 4°C or at 150 rpm in a crystallizer (*Crystal16*, Technobis) at 25°C. For all investigated mRNA sequences, a significant reduction of precipitate aggregates in the supernatant was observed after a short centrifugation period at 12000 rpm, in a *Centrifuge 5425 R* by Eppendorf, (see Figure 1c) where, after 15 min, no aggregates were observed in any of the investigated sample compositions. However, after leaving the samples for about 5 to 10 min, a resuspension of the aggregates was observed, likely due to diminishing density differences between the solid and liquid phase from the incorporation of water into the aggregates. Thus, centrifuging the equilibrated samples for 15 min at 12000 rpm and their respective equilibration temperature with rapid subsequent sampling was deemed sufficient for this study. The result is a neat separation of the supernatant and precipitate phases (shown visually in Figure 1a, right illustration). More information about these experiments is given in the SI of this work. The liquid phase was sampled and its concentration quantified via a Thermo Fischer Scientific *NanoDrop One*^*c*^ UV-Vis spectrophotometer based on absorbance at wavelengths of 260 and 280 nm. All samples were prepared and measured in triplicate.

**Figure 1:**
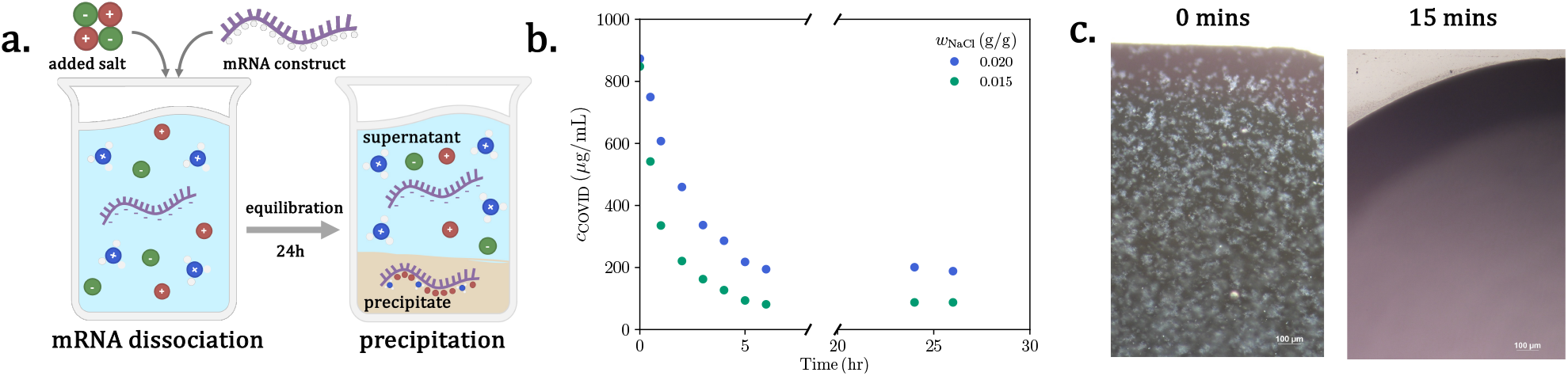
**a.** Precipitation procedure where neat mRNA sequences and salts are mixed in water, totally ionizing the mRNA chain. The mRNA chain then complexes with the added cation and hydronium to form a precipitate which separates from the supertant after equilibration over 24 hr. **b**. Concentration change of COVID sequence over equilibration period. **c**. Microscopic images of droplets sampled from aqueous mRNA/precipitant/salt supernatent after varying centrifuging times at 12000 rpm.

### SAFT-*γ* Mie Equation

To develop a framework capable of predicting the solubility of mRNA sequences under a variety of conditions (e.g., solvent, temperature, salt concentration), it is first necessary to consider the types of interactions present in such systems. As illustrated in Figure 2a, an mRNA strand is composed of four nucleobases (uracil, cytosine, guanine, and adenine), deoxyribose, and phosphate groups. Together, these substructures give rise to nearly all major classes of molecular interactions. Aromatic and cycloalkyl groups contribute hydrophobic (dispersive) interactions; hydroxyl, ketone, and nitrogen-containing groups enable hydrophilic (associative) interactions; and the phosphate backbone provides electrostatic interactions. These act simultaneously and often competitively. Even within a single interaction class, the chemical specificity of each functional group introduces subtle differences in both self-interactions and cross-interactions. Additional complexity arises from solvent molecules (e.g., water, ethanol, PEG) and inorganic salts (e.g., sodium chloride, sodium acetate, ammonium acetate), which further complicate the interaction landscape.

**Figure 2:**
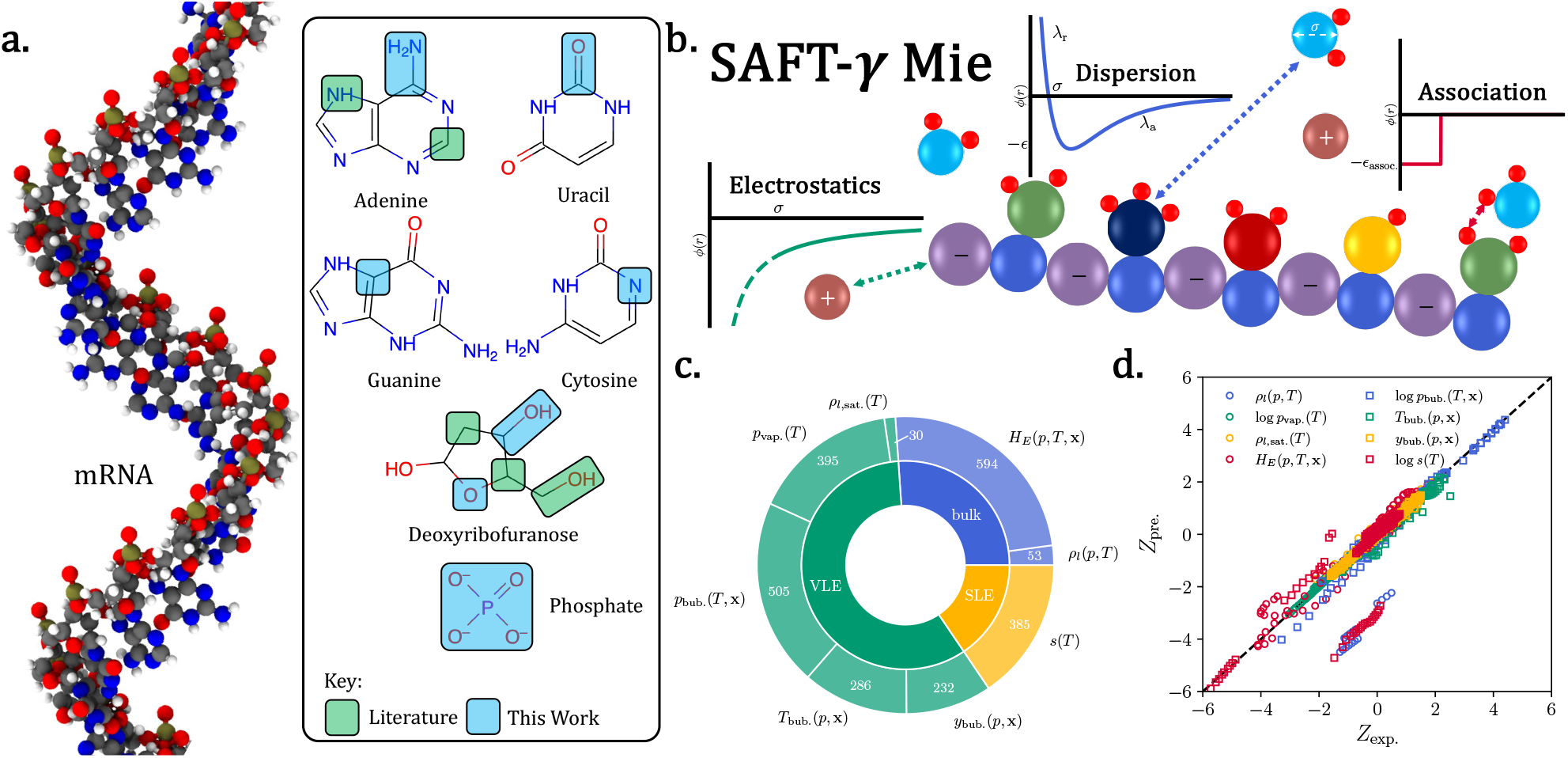
Parameterization of SAFT-*γ* Mie for mRNA modeling. **a.** Fragmentation of an mRNA strand into its monomers (nucleobases, deoxyribofuranose and phosphate) and further reduction to the constituent functional groups (see Model Parameterization for explanation). **b**. Simplified representation of molecules within the SAFT-*γ*-Mie equation, accounting for dispersive, association and electrostatic interactions. **c**. Type and quantity of data included within the parameterization. **d**. Parity plot comparing the SAFT-*γ* Mie predictions with experimental data considered within the parameterization.

Accordingly, a model is needed that can represent the chemical complexity of mRNA-containing systems without depending heavily on system-specific experimental input. Here, we adopt a physically derived approach: the statistical associating fluid theory (SAFT) [24, 25]. Specifically, we employ the *γ*-Mie variant [26–28], which represents molecules as hetero-segmented chains interacting through the relevant physical forces (illustrated in Figure 2b). This approach provides an analytical expression for the Helmholtz free energy, *A*, in terms of the system volume (*V*), temperature (*T*) and composition (**n**) as the sum of six contributions:

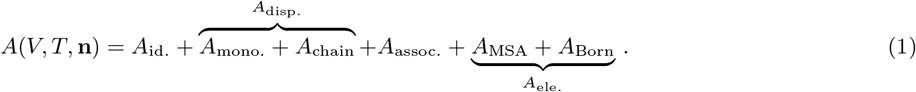

The derivation of (1) can be found elsewhere [25, 29–31]. Below, we briefly summarize the role of each contribution.

The *A*_id._ term corresponds to the ideal contribution, capturing translational degrees of freedom and serving as the reference state. Rotational and vibrational kinetic terms, while relevant for thermal properties, do not affect equilibrium solubility and are therefore neglected. The *A*_mono._ term describes monomer-level Mie interactions between disconnected functional groups (shown visually by the blue curve in Figure 2b):

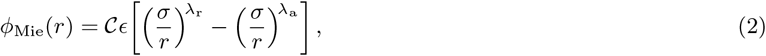

where *r* is the inter-monomer distance, *ϵ* the well depth, *σ* the monomer diameter, and *λ*_r_ and *λ*_a_ the repulsive and attractive exponents, respectively. The prefactor *C* ensures the minimum of the potential is ^*−*^*ϵ*, for any particular choice of *λ*_r_ and *λ*_a_. The interactions between pairs of functional group is characterized by a unique parameter set, reflecting chemical heterogeneity (denoted by the distinct colors in Figure 2b). The *A*_chain_ term then accounts for the entropy reduction upon connecting monomers into chains. Together, *A*_mono._ and *A*_chain_ form the dispersion contribution (*A*_disp._), which effectively captures van der Waals–type, hydrophobic interactions.

The *A*_assoc._ term represents directional association interactions, modeled using a square-well potential (red curve in Figure 2b):

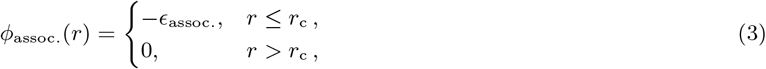

where *ϵ*_assoc._ is the association well depth and *r*_c_ the cutoff radius. In practice, a bonding volume (*K*_bond_) replaces *r*_*c*_ and can be thought of as a measure of steric hindrance. Each associating pair is thus parameterized by *ϵ*_assoc._ and *K*_bond_. Unlike dispersion forces, association interactions are short-range, strong, and highly directional, designed to emulate hydrogen bonding. Within mRNA, they arise between the various electron donor and acceptor groups. The flexibility of this contribution allows SAFT-*γ* Mie to more rigorously capture complex interactions in mRNA systems, such as water hydration of phosphate groups and chain self-interactions. However, a key limitation is the assumption that these interactions are treated in an average manner, both spatially and conformationally, rather than being explicitly conformer-specific. As a result, SAFT-*γ* Mie does not explicitly describe secondary structures, although the association term partially accounts for the additional stability arising from self-interactions between nucleobases.

Electrostatic interactions, characterized by the Coulombic potential:

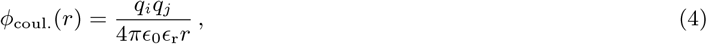

where *q*_*i*_ and *q*_*j*_ are the charges of interacting species, *ϵ*_0_ is the vacuum permittivity, and *ϵ*_r_ is the relative permittivity, are included through the *A*_MSA_ term, derived via the mean spherical approximation [32, 33]. The mean-spherical approximation has been successfully applied to the modeling of complex systems such as polyelectrolytes, of which mRNA may be considered a subclass, yielding a good degree of accuracy [34–36]. Within SAFT-*γ* Mie, it is assumed that *ϵ*_r_ depends on the system temperature, volume and composition of the *solvents* [37]. The effect of charged species on *ϵ*_r_ has been neglected primarily for simplicity, but also because of the difficulties associated with including the influence of charged species on the relative permittivity [38]. Finally, as mixed solvent conditions will be explored in this work, the change in solvation environment is accounted for through the inclusion of the Born term, *A*_Born_. The combination of these two terms will be referred henceforth as the electrostatic contributions.

The full SAFT-*γ* Mie implementation is available through the open-source package Clapeyron.jl [39].

### Model Parameterization

As outlined above, a number of parameters must be determined to fully characterize the interactions in an mRNA-containing system. Although these parameters are physically motivated and can, in principle, be obtained from *ab initio* calculations [40, 41], they are more commonly estimated by regressing the SAFT-*γ* Mie parameters against experimental data. Direct regression on the target mRNA sequence is generally infeasible due to the scarcity of solubility and thermophysical data. Moreover, parameters fitted to a single sequence are unlikely to be transferable to other sequences. A more robust strategy is to parameterize constituent substructures (nucleobases, nucleosides, and nucleotides) so that the resulting parameters can be transferred to different sequences following the principles of group-contribution approaches. Yet, even at the substructure level, experimental data remain limited. For this reason, we focus instead on the functional groups present within these substructures (highlighted in Figure 2a). Experimental measurements are considerably more abundant for small molecules containing one or several of these groups. Parameters fitted to such simpler species should, in principle, remain transferable to larger biomolecular systems. Conveniently, many of these functional groups (highlighted in green in Figure 2a) have already been parameterized in previous SAFT-*γ* Mie studies [28], leaving only a subset (highlighted in light blue) requiring new parameter estimation.

To adequately represent functional group interactions, we assembled a diverse experimental dataset spanning more than 40 species and approximately 2000 data points. This dataset includes bulk properties (liquid densities, *ρ*_*l*_, and excess enthalpies, *H*_*E*_), vapor–liquid equilibrium properties (saturated liquid densities, *ρ*_*l*,sat._, saturation pressures, *p*_sat._, bubble pressures, *p*_bub._, bubble temperatures, *T*_bub._, and vapor-phase compositions at the bubble point, *y*_bub._), and solid–liquid equilibrium solubilities (*s*). The methods used to estimate these properties using SAFT-*γ* Mie are omitted here as we follow established estimation methods already implemented in Clapeyron.jl [39]. The size and distribution of the dataset are summarized in Figure 2c Although modest compared to datasets used in purely data-driven approaches [42], this collection is near comprehensive with respect to available measurements for the relevant functional groups. Furthermore, it significantly exceeds the datasets typically used to fit SAFT-*γ* Mie parameters [26]. Importantly, the physically grounded nature of SAFT-*γ* Mie inherently reduces the need for large datasets, provided the selected data adequately capture the relevant range of interactions. For this reason, the dataset is intentionally biased toward vapor–liquid equilibrium properties rather than the target property of solid–liquid equilibrium solubilities. The vapor–liquid transition provides especially rich information on intermolecular interactions, since it spans the regime from dense liquid packing, where these interactions are most pronounced, to the near-ideal behavior of the vapor phase. We therefore consider the assembled dataset sufficient for robust parameterization. Complete references for the experimental data subsets are provided in the Supporting Information.

Parameter fitting was carried out using the Metaheuristics.jl [43] package to perform global optimization. The weighted least-squares objective function

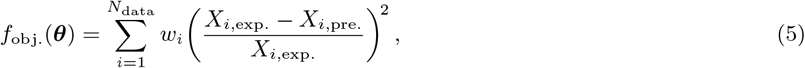

was minimized, where ***θ*** denotes the vector of parameters being optimized, *w*_*i*_ is a weighting coefficient empirically adjusted to reflect the reliability of the experimental measurement, *X*_*i*,exp._ is the experimental value of the *i*th data point, and *X*_*i*,pre._ is the corresponding SAFT-*γ* Mie prediction.

### Model Performance

The performance of the fitted parameters is summarized by the parity plot in Figure 2d. To allow direct comparison across different types of properties, all data were normalized using the *z*-score; for properties spanning several orders of magnitude, logarithmic scaling was applied prior to normalization. As shown, most of the experimental data fall on or near the parity line, demonstrating the high accuracy of the model across diverse systems, properties, and conditions. Only two datasets, one solubility and one liquid density, show significant deviation. These correspond to either chemically complex systems or cases where experimental uncertainty led to the assignment of low weighting coefficients.

Following the large-scale regression, additional refinement of parameters (particularly those with limited experimental data) was performed using a focused dataset of aqueous solubilities for nucleobases and nucleosides. This step improved transferability of the parameters to the substructures most relevant for mRNA. Details of these refinements are provided in the Supporting Information. For the few remaining parameters lacking experimental data, combining rules [38, 44] were employed.

Although this parameterization strategy is more demanding than generalized data-driven approaches, its rigor provides a key advantage: parameter values can be continuously assessed for physical plausibility. Such validation can be carried out by comparison to *ab initio* calculations or to values reported in the literature. For example, the association interactions between nucleobase hydrogen-bonding sites are found to be significantly stronger than other association interactions, consistent with chemical intuition. This level of interpretability stands in contrast to the “black-box” character of many machine-learning–based fitting procedures. The final parameter set, along with notes on any special treatment of specific interactions, is provided in the Supporting Information.

While developed here for mRNA systems, the resulting group-contribution parameters are inherently transferable. By extension, they may also be applied to other biomolecules, pharmaceutical compounds, and polyelectrolytes, extending the utility of this work beyond the immediate target system.

### Thermodynamic Modeling of mRNA Precipitation

As illustrated in Figure 1a, when mRNA is dissolved in aqueous solution, the phosphate backbone is expected to be fully ionized due to its very low pK_a_. In the absence of added salts, hydronium ions serve as the natural counterions. This assumption is consistent with the high solubility of mRNA in pure water, where precipitation is not expected. Upon addition of salt, which dissociates into cations and anions, the system can be considered to contain five species (mRNA, hydronium, cation, anion, and water), six in the case where a co-solvent is added.

At equilibrium, the system is assumed to consist of two phases: a liquid phase (the supernatant), containing all species, and a solid phase (the precipitate), composed of mRNA complexed with condensed cations. The conventional assumption is that the cations in the precipitate correspond solely to the salt cations (e.g., sodium or ammonium) [45]. However, given the strong hydrogen-bonding interactions between hydronium and the electron-donating groups of mRNA (particularly the phosphate groups), we postulate that the precipitate contains both salt cations and hydronium ions coordinated to the mRNA backbone. The complexation with hydronium can also be interpreted as accounting for the present of water in the mRNA precipitate. The equilibrium solubility of mRNA is determined by enforcing chemical equilibrium between the solid and liquid phases of the mRNA–cation complex (*µ*_mRNA*†*_):

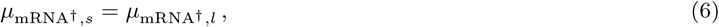

where *µ*_*i,α*_ is the chemical potential of species *i* in phase *α*. In this case, the mRNA-cation complex (mRNA^†^) is made up of the mRNA sequence, as well as the counterions. For the liquid phase, the chemical potential of the complex is expressed as

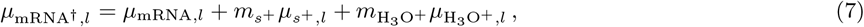

where *m*_*i*_ denotes the number of species *i* associated with each mRNA strand, sufficient to neutralize its charge. Each chemical potential on the right-hand side of Eq. (7) is obtained from the Helmholtz free energy (Eq. (1)) by differentiation with respect to species composition:

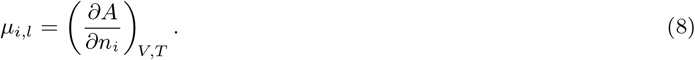

Modeling the solid phase presents additional challenges, as SAFT-*γ* Mie is inherently restricted to fluid-phase properties and cannot directly describe solid states. To address this, we adopt a reference-state approach, whereby the chemical potential of the solid phase is expressed relative to that of the charged species at infinite dilution in water. Subtracting this reference, 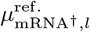from both sides of Eq. (6) yields

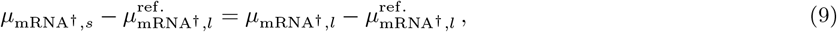

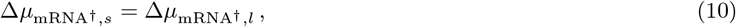

where 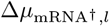 is obtained by subtracting the reference state from each term in Eq. (7). Using the Helmholtz–Gibbs equation, 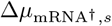 can be obtained as

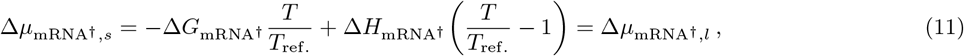

where Δ*G*_mRNA*†*_ and Δ*H*_mRNA*†*_ are the Gibbs free energy and enthalpy of dissolution of the mRNA precipitate at infinite dilution, evaluated at a reference temperature *T*_ref._ (typically set to 298.15 K). For simple salts, these quantities are typically available in the literature. In contrast, for mRNA, they must be estimated by fitting to experimental solubility data, along with the stoichiometric number of condensed salt cations (*m*_*s*+_) per mRNA strand. Because these parameters depend on the identity of the salt cation, two parameter sets are required: one for sodium and one for ammonium. We further assume that Δ*G*_mRNA*†*_, Δ*H*_mRNA*†*_, and *m*_*s*+_ scale proportionally with the number of base pairs in the mRNA sequence. In this work, we fit parameters using aqueous solubility data for the FLuc sequence and validate model transferability by extrapolation to the COVID sequence. Through rearrangement, the mRNA concentration, *c*_mRNA_, can be expressed from (11) as

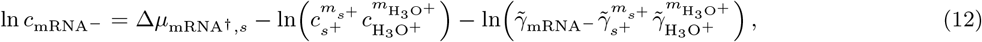

Where 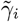 denotes the molality-based activity coefficient of species *i*, referenced to infinite dilution in water. This framework is also applicable to mixed-solvent systems, as the precipitate and reference state remain unchanged upon introduction of additional solvents.

## Results and Discussions

### Aqueous Solubility of mRNA sequences

The experimental and theoretical solubilities of the two mRNA sequences are shown in Figures 3a and 3b. In both cases, solubility decreases monotonically with added salt, consistent with mRNA–cation complexation that promotes precipitation. This effect is further enhanced at lower temperatures, where unfavorable energetic contributions (dispersion and association) dominate over entropic contributions that would otherwise favor mixing.

**Figure 3:**
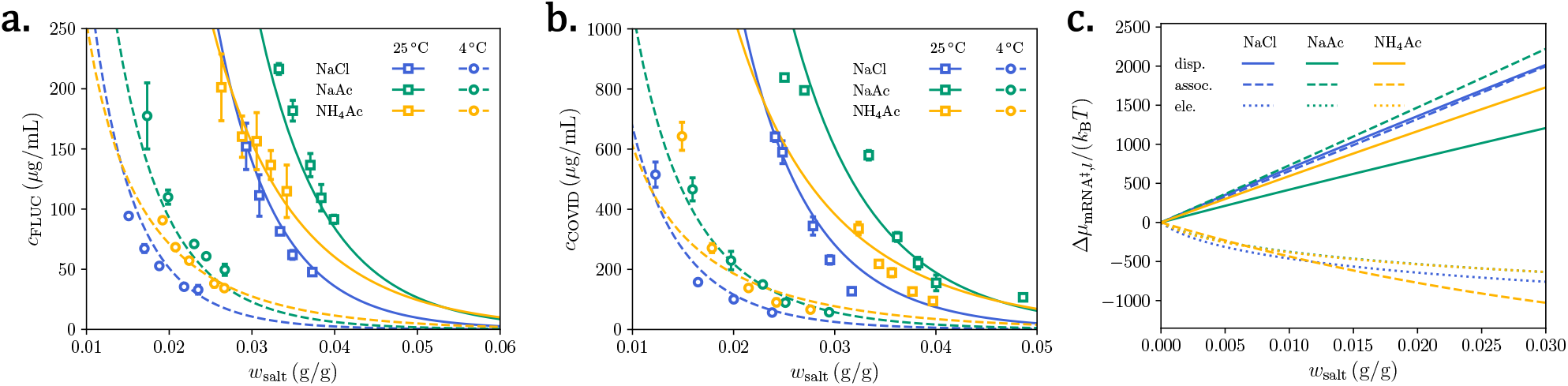
Precipitation of mRNA sequences in an aqueous environment with added salt and varying temperature. **a.** FLuc and **b**. COVID sequence solubility in water with added salt (weight fraction, *w*_salt_) where symbols denote experimental data and lines represent estimates from SAFT-*γ* Mie. The latter are predictions from the theory. **c**. Contribution to the excess chemical potential of the mRNA complex in water with addition of various salts. The reference system (denoted by *‡*) is taken to be an aqueous solution of mRNA at a composition of *w*_mRNA_ = 0.01.

Direct comparison across salts reveals only modest quantitative differences. Replacing chloride with a more kosmotropic ion [46], acetate (NaCl *→* NaAc), slightly increases solubility, as expected. On the other hand, replacing sodium with ammonium (NaAc *→* NH_4_Ac), which is also more kosmotropic, actually decreases solubility. This can be somewhat rationalized by the fact that when oppositely charged kosmotropic ions are introduced in solution, favorable interactions between them reduce the kosmotropic effect [47]. Interestingly, while NaCl and NaAc exhibit similar slopes, NH_4_Ac shows a weaker concentration dependence. These trends suggest subtle ion-specific effects, possibly arising from donor–acceptor interactions between acetate or ammonium and functional groups in mRNA, though the precise driving forces are difficult to identify unambiguously.

The fitted SAFT-*γ* Mie results (parameters provided in the supplementary material) show excellent agreement with experiment (Figure 3a). The model successfully reproduces quantitative salt-specific differences at both temperatures, most clearly in the case of NaCl and NaAc, where the precipitated complex shares the same cation. In this case, differences arise solely from the SAFT-*γ* Mie theory, thus highlighting its ability to distinguish ions along the Hofmeister series. Remarkably, when extrapolated to the COVID sequence for validation (Figure 3b), the model also yields qualitatively accurate predictions simply by scaling precipitate parameters with the number of base pairs. The model continues to capture salt-specific differences, with residual quantitative deviations likely reflecting differences in RNA purity between the FLuc and COVID sequences.

With a reliable predictive framework established, we next examined the physical origins of salt-driven precipitation by decomposing the mRNA chemical potential (Eq. (7)) into dispersion, association, and electrostatic contributions in the liquid phase. Each contribution was normalized by subtracting a reference chemical potential for an aqueous mRNA solution at *w*_salt_ = 0 and *w*_mRNA_ = 0.01, as shown in Figure 3c. Positive contributions 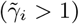correspond to unfavorable (demixing) interactions, while negative contributions 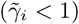indicate favorable (mixing) ones.

For all salts, the electrostatic contribution is favorable and becomes increasingly so at higher salt concentrations. This is expected [32, 33, 35, 38], as added salt increases ionic strength and shortens the screening length, thereby enhancing solubility. However, when discussing the role of electrostatics in polyelectrolyte precipitation, conventional arguments invoke counterion condensation, where cations bind directly to phosphate backbone, as the main driving force. While theoretical approaches accounting for counterion condensation do exist, the role of counterion condensation is still debated within literature [48, 49]. Within SAFT-*γ* Mie, counterion condensation is not explicitly included within the electrostatic term. Differences between salts in this term primarily reflect differences in ion size: acetate, being larger than chloride, produces a slightly weaker electrostatic interaction.

Dispersive interactions show a noticeably different trend. Inorganic ions such as Na^+^ and Cl^*−*^ have very weak dispersive forces, resulting in unfavorable interactions with the more-polarizable mRNA functional groups. On the other hand, more polarizable ions like acetate and ammonium will interact more favorably. We note that, because experimental data are sparse, cross-interaction parameters for these ions are estimated using combining rules, though the resulting trends remain physically reasonable.

Association interactions provide the largest overall contribution. For NaCl and NaAc, the effect is strongly unfavorable, reflecting the disruptive role of ions on hydrogen bonding between mRNA and itself, and with water. The effect is particularly pronounced for acetate, which itself associates with water, thereby reducing water’s availability to interact with mRNA sites. It is interesting to note that, while such an effect is expected from a kosmotropic ion, it is surprising that this effect contributes unfavorably to the solubility of mRNA. It highlights that it is actually the more polarizable nature of kosmotropic ions that contributes to the improved solubility of mRNA. In contrast, NH_4_Ac exhibits an unexpected favorable association contribution. Although ammonium does associate with water (as expected from a kosmotropic ion), it can also associate strongly with phosphate groups. The latter interaction dominates, leading to an overall favorable interaction. While the combination of dispersive and association interactions seem to indicate that NH_4_Ac should improve the solubility of mRNA, differences in the solid-phase chemical potential lead to the overall reduction in solublity. Nonetheless, these interactions likely underline the distinctive slope of the NH_4_Ac solubility curve observed in Figures 3a and 3b.

### Mixed Solvent Solubility of mRNA sequences

Two solvents were considered in this work, both of which are commonly employed in mRNA precipitation: ethanol and PEG6k. Experimental data were collected for both solvents at five compositions, in combination with all salts discussed previously. Because the overall trends were similar, only the sodium chloride results are presented in Figure 4; the remainder are available in the supplementary information. The experimental procedure was unchanged, though, as shown in Figure 4c, in contrast to the aqueous case (Figure 3b), the mRNA concentration was observed to oscillate around an equilibrium value rather than monotonically approach a certain value. This behavior introduced additional uncertainty relative to the aqueous measurements.

**Figure 4:**
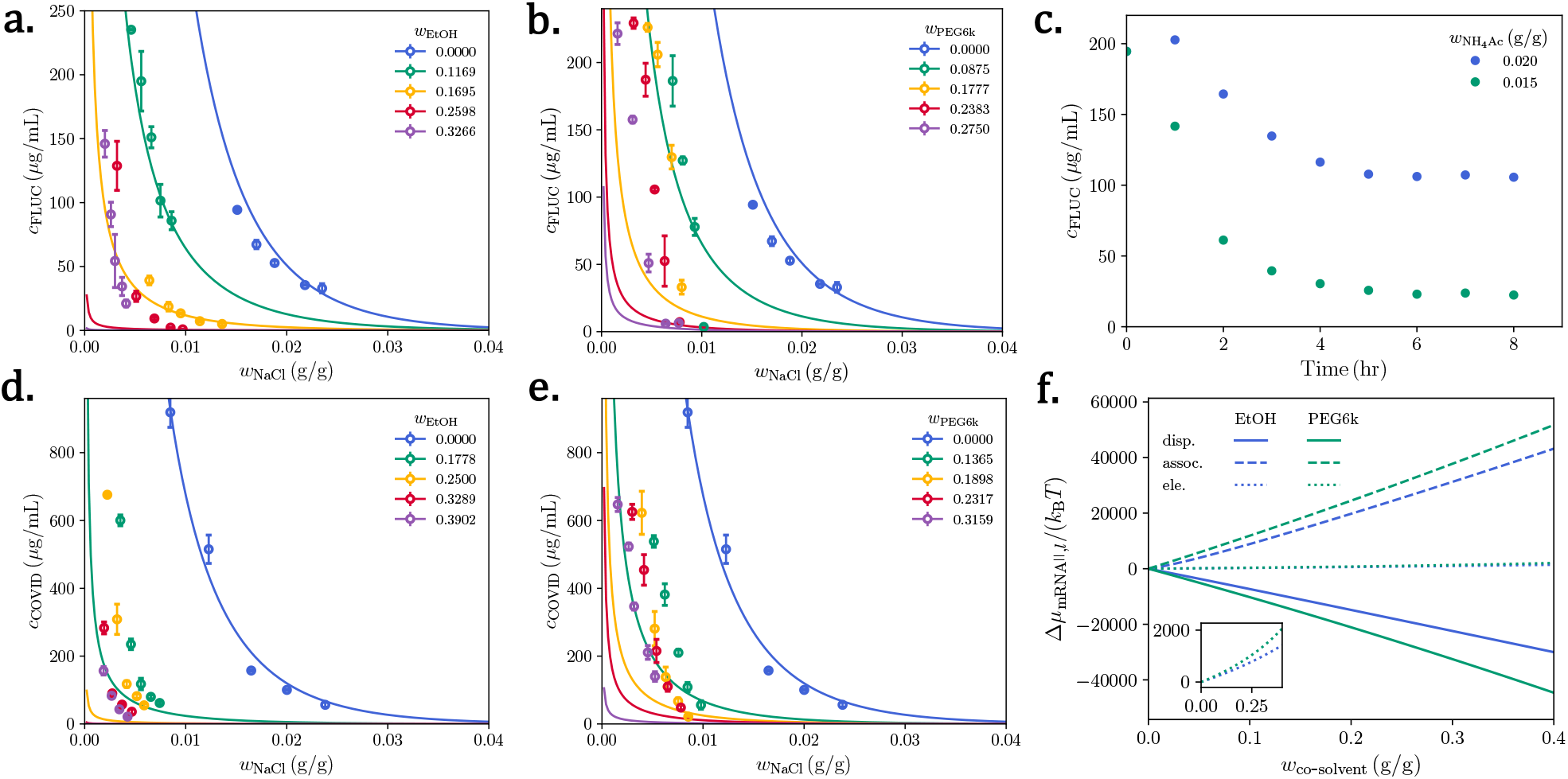
Precipitation of mRNA sequences in a mixed solvent environment with added salt at 4°C. **a**.+**b.** FLuc and **d**.+**e**. COVID solubility in water+ethanol (**a**.+**d**.) and water+PEG6k (**b**.+**e**.) solutions with added sodium chloride (weight fraction, *w*_NaCl_) where symbols denote experimental data and lines represent estimates from SAFT-*γ* Mie. **c**. Concentration change of FLuc sequence in a water+ethanol solution over equilibration period. **f**. Contribution to the excess chemical potential of the mRNA complex in a mixed solvent with varying co-solvent mass fraction. The reference system (denoted by the || superscript) is taken to be an aqueous solution of mRNA at a composition of *w*_mRNA_ = 0.01 and NaCl composition of *w*_NaCl_ = 0.015.

The experimental results in Figures 4abde show that the addition of either ethanol or PEG6k significantly decreases mRNA solubility. With respect to the solvation free energy, since PEG6k has a lower relative permittivity than ethanol [50], it acts as a slightly more effective precipitant. We also find that the COVID sequence is more strongly affected by co-solvent addition than FLuc, likely because of its larger size, which makes it more sensitive to perturbations in a solvation environment.

The SAFT-*γ* Mie predictions shown in Figure 4 represent total predictions with no additional fitting, as the parameters obtained for aqueous systems are assumed to transfer directly to mixed-solvent systems. For ethanol and PEG6k, the model performs remarkably well, achieving near-quantitative agreement at moderate solvent concentrations. At higher co-solvent fractions, deviations are more pronounced, though the predictions remain physically sensible. This discrepancy may reflect changes in the composition of the mRNA–cation complex as water is displaced by co-solvent, an effect not explicitly captured in the current model since it would require additional parameters. For the COVID sequence (Figures 4d and 4e), agreement is primarily qualitative, particularly at high co-solvent fractions, consistent with the poorer performance already observed in aqueous systems. Additional experimental uncertainty and lower sample purity further contribute to the discrepancies.

To probe the molecular origin of solvent effects, the SAFT-*γ* Mie contributions were decomposed relative to a reference aqueous solution containing *w*_mRNA_ = 0.010 and *w*_NaCl_ = 0.015 (Figure 4f). The total electrostatic contribution decreases slightly with co-solvent addition, as expected from the reduction in relative permittivity, and is more unfavorable for PEG6k than ethanol. However, this effect is small compared to other contributions, since the dielectric constant changes only modestly at the co-solvent compositions considered here. At higher co-solvent contents, where the dielectric decrement is larger, the electrostatic contribution would be expected to play a more significant role.

Dispersion interactions provide a much stronger contribution. For both ethanol and PEG6k, these interactions become favorable because the alkyl chains of the co-solvents interact with the aromatic and cycloalkyl groups of mRNA, stabilizing the system. Importantly, PEG6k, due to its polymeric nature and extended hydrophobic backbone, introduces a larger dispersive stabilization than ethanol, though this is counterbalanced by its stronger disruption of association interactions (see below). These results suggest that dispersive effects represent an important lever for tuning mRNA precipitation, particularly when co-solvents contain large, flexible, or highly polarizable groups.

Association interactions dominate the overall solvent response. Both ethanol and PEG6k disrupt hydrogen bonding between water and mRNA by competing for the same donor and acceptor sites. Ethanol, with its single hydroxyl group, competes primarily through hydrogen bonding with water, thereby weakening hydration of phosphate and nucleobase sites. PEG6k, by contrast, contains multiple ether oxygens along its backbone that can act as electron donors. This creates many competing association sites, dramatically reducing the number of hydrogen bonds between water and mRNA. The result is a much stronger unfavorable association contribution for PEG6k than ethanol, which directly explains why PEG6k is the more effective precipitant despite its larger dispersive stabilization.

### Extrapolation to other conditions

In the previous sections, we focused on conditions for which experimental data were available. However, a key advantage of an approach such as SAFT-*γ* Mie is its ability to extrapolate to conditions beyond those directly measured. This was already demonstrated in the case of mixed solvents, but the framework is general and can be extended along multiple dimensions, as illustrated in Figure 5.

**Figure 5:**
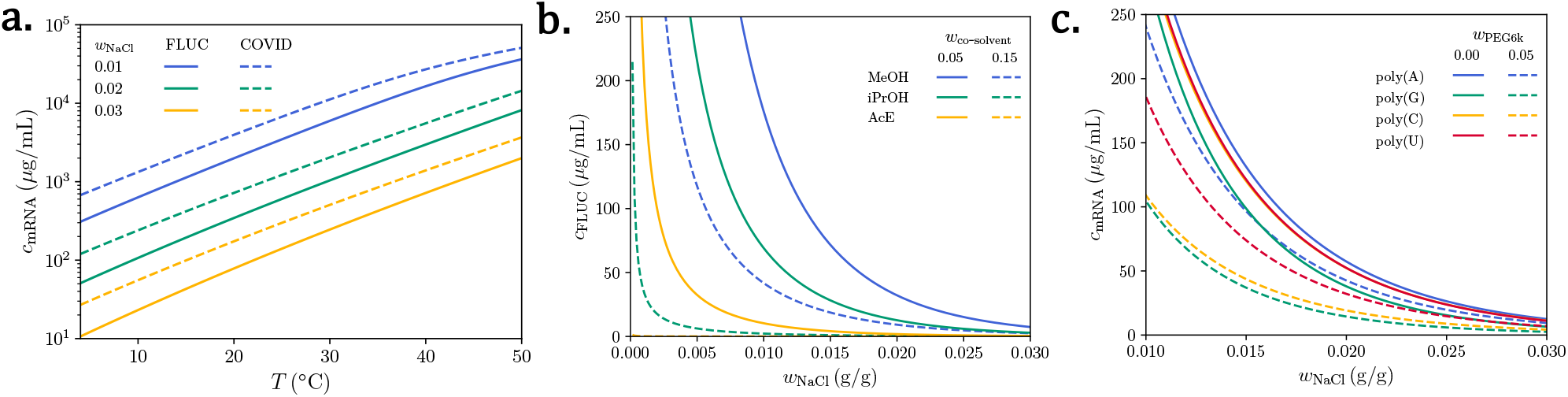
mRNA solubility predictions using SAFT-*γ* Mie. **a.** Aqueous solubility of FLuc and COVID with varying NaCl concentration, extrapolating to higher temperatures. **b**. Solubility of FLuc with added salt using unseen co-solvents (MeOH: methanol, iPrOH: isopropanol, AcE: acetone). **c**. Solubility of various homonucleotides with varying NaCl concentration and added PEG6k.

Figure 5a shows extrapolation of the two mRNA sequences to higher temperatures at varying salt concentrations. Consistent with Figures 3a and 3b, mRNA solubility increases with temperature, in line with the functional form of the solid-phase chemical potential. At high temperatures, particularly for the lowest salt concentrations in COVID, small deviations in the trends arise. These deviations arise from non-idealities within SAFT-*γ* Mie, where unfavorable enthalpic contributions from dispersion or association begin to dominate.

While the previous section focused only on ethanol and PEG6k as co-solvents, SAFT-*γ* Mie is equipped with a broad parameter database that allows modeling of many additional co-solvents. In Figure 5b, we examine three less common precipitants: methanol, isopropanol, and acetone. Methanol and isopropanol, like ethanol, both possess hydroxyl groups and yield higher solubilities than acetone. The bulkier nature of isopropanol likely leads to greater disruption of association interactions compared to methanol, resulting in slightly lower solubility. In contrast, acetone, being an aprotic solvent, primarily disrupts water–mRNA hydrogen bonding without offering compensating associations, thereby strongly promoting precipitation. At higher co-solvent fractions, acetone drives near-complete precipitation of mRNA. These trends coincidentally follow the relative dielectric constants of the solvents, in agreement with physical intuition.

A natural question concerns the transferability of the fitted parameters, particularly Δ*G*_mRNA*†*_ and Δ*H*_mRNA*†*_, to other mRNA sequences. Both FLuc and COVID strands studied here have relatively uniform nucleobase distributions, and we therefore expect the parameters to extrapolate reasonably to other disordered sequences. While SAFT-*γ* Mie is expected to perform well for single-stranded mRNA sequences that remain relatively disordered, its accuracy may degrade for highly ordered systems that adopt stable, sequence-specific conformations, such as large diblock or structured constructs (e.g., poly(A–co–U)). In these cases, precipitation behavior is dominated by conformational effects not captured within the mean-field nature of the theory. To test the model’s transferability under more ideal conditions, Figure 5c presents predictions for four homogeneous single-stranded sequences of equal length at two PEG6k fractions and varying NaCl concentrations. The predicted solubilities are both physically reasonable and consistent with experimental expectations: poly(adenylic acid) *>* poly(methyl-pseudouridylic acid) *>* poly(cytidylic acid) *>* poly(guanylic acid). This trend reflects the greater propensity of cytosine- and guanine-containing chains to self-aggregate (e.g., via G-quadruplex formation), in contrast to adenine- and uracil-containing strands. Although SAFT-*γ* Mie does not explicitly account for such secondary structures, these effects are likely represented implicitly through the fitted parameters. Since the solid-phase parameters were held constant, the observed differences arise solely from the SAFT-*γ* Mie free-energy contributions, demonstrating the model’s ability to capture nucleobase-specific effects and extrapolate across different single-stranded sequences. The relatively small differences in solubility between homopolymers may appear surprising, given that the isolated nucleobases differ in solubility by orders of magnitude (see SI)[51, 52]. This can be rationalized by noting that, as shown in Figures 3f and 4c, the dominant factor governing precipitation is the solvation of the phosphate groups. Because all chains contain the same number of phosphates, their overall solubilities are expected to be similar, with nucleobase identity contributing only a secondary correction. These effects become more pronounced in the presence of PEG6k, where unfavorable interactions between the nucleobases and the co-solvent further modulate solubility.

These results highlight not only the quantitative predictive power of SAFT-*γ* Mie across temperature, solvent, and sequence variations, but also its ability to yield physically interpretable insights into the molecular interactions that govern mRNA precipitation.

## Conclusions

In this study, we have developed and validated the SAFT-*γ* Mie framework for predicting the precipitation behavior of mRNA across a wide range of conditions. By parameterizing at the level of functional groups, rather than individual sequences, we achieved both accuracy and transferability, allowing the model to reproduce experimental solubilities for different salts, solvents, and temperatures with remarkable agreement. The rigorous parameterization strategy, though more demanding than generalized data-driven approaches, ensures physical plausibility at every stage and avoids the black-box character often associated with machine learning–based fitting.

Beyond predictive accuracy, the framework provides mechanistic insight into the molecular origins of mRNA precipitation. Analysis of the Helmholtz free energy contributions reveals that, while electrostatics and dispersive stabilization play important roles, it is the disruption of water–mRNA hydrogen bonding that primarily drives precipitation. This physical interpretation is consistent with experimental observations and clarifies why certain salts and solvents act as stronger precipitants. Moreover, the model captures sequence-dependent solubility trends without additional fitting, highlighting its ability to extrapolate to unseen sequences and providing a first-principles explanation for differences between nucleobase types.

Taken together, these results demonstrate that SAFT-*γ* Mie offers a powerful combination of predictive accuracy, transferability, and interpretability. The framework not only reproduces experimental data under measured conditions but also extends confidently to mixed solvents, elevated temperatures, and different sequences. As such, it provides both a reliable predictive tool and a physically transparent framework for guiding the rational design of mRNA formulations and precipitation processes.

## Supplementary Material

We provide the following supplementary material:

- si.pdf: Supplementary information providing additional details regarding the experimental procedure, discussion regarding the parameterization of SAFT-*γ* Mie and supplementary figures.
- params.zip: Zip file containing all parameters used within the SAFT-*γ* Mie model.
- data.zip: Zip file containing all data collected from experiments in this work.

## Data Availability Statement

The experimental data, implementation of the thermodynamic model, and example codes for deploying the model can be found at https://github.com/pw0908/mRNA_thermodynamic_modelling, as well as at Zenodo: https://doi.org/10.5281/zenodo.17388262.

## Author Contribution Statement

Vico Tenberg: Conceptualization (Experimental), Data Curation, Investigation, Methodology (Experimental), Visualization, Writing - Original Draft Preparation. Pierre J. Walker: Conceptualization (Theory and Modeling), Data Curation, Formal Analysis, Investigation, Methodology (Theory and Modeling), Software, Visualization, Writing - Original Draft Preparation. Pavan K. Inguva: Conceptualization (Theory and Modeling), Investigation, Methodology (Theory and Modeling), Writing - Original Draft Preparation. Maria del Carme Pons Royo: Investigation, Methodology (Experimental). Andrew L. Acevedo: Investigation. Vernon S. Lin: Investigation. Marion Pang: Writing - Review & Editing. Zhen-Gang Wang: Writing - Review & Editing. Allan S. Myerson: Funding Acquisition, Project Administration, Supervision, Writing - Review & Editing. Richard D. Braatz: Funding Acquisition, Project Administration, Supervision, Writing - Review & Editing.

## Acknowledgments

This research was supported by the U.S. Food and Drug Administration under the FDA BAA-22-00123 program, Award Number 75F40122C00200. Financial support is also acknowledged from the Agency for Science, Technology and Research (A*STAR), Singapore.

