## Supplementary Material for "Thermodynamic Modeling of mRNA with the Addition of Precipitants"

#### Experimental Methods

##### Equilibrium Studies

In preliminary experiments, prior to the solubility determination, equilibrium studies for each of the regarded mRNA constructs were performed. For this, various supersaturated mixtures were created and equilibrated at 4°C. For these equilibration experiments, a liquid phase sample was taken each hour and its concentration measured until a stable value was reached. Before the sampling, the suspension was centrifuged and shaken after the sampling to allow for resuspension of the aggregates. For all investigated constructs and compositions, an equilibration period of about 6 h was observed. Additionally, in equilibrium, stable values without degradation of the mRNA were observed for at least 48 h. Thus, for the solubility determination, an equilibration time of 24 h was chosen.

##### Centrifugation of mRNA Suspensions

To verify sufficient solid-liquid separation during the centrifugation, several samples of varying composition and of each mRNA construct were prepared. Their supernatent solution was sampled after being centrifuged at 12000 rpm for certain durations. The liquid phase was then placed on a microscope slide and observed under a microscope. Selected pictures and corresponding centrifugation times are shown in Figure SI.1.

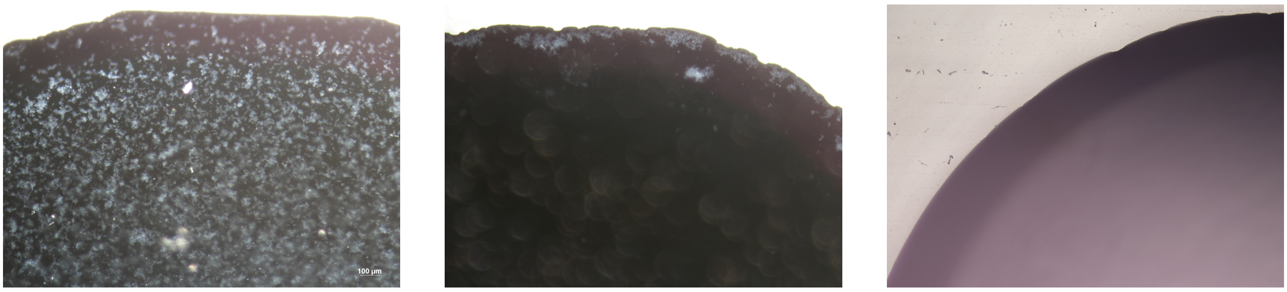


(a) 0 min    (b) 2 min    (c) 15 min

Figure SI.1: Selected microscopic images of droplets sampled from aqueous mRNA/precipitant/salt supernatent after varying centrifuging times at 12000 rpm.

For all investigated mRNA constructs, a significant reduction of precipitate aggregates in the supernatent was observed after a short centrifugation period (see Figure SI.1 (a) and (b)). After 15 min, no aggregates were observed in any of the investigated sample compositions. However, after leaving the samples for about 5 to 10 min, a resuspension of the aggregates was observed, likely due to diminishing density differences between the solid and liquid phase due to incorporation of solvent into the aggregates. Thus, centrifuging the equilibrated samples for 15 rpm with rapid subsequent sampling was deemed sufficient for this study.

### Theory

A number of assumptions were made with respect to cross-interaction parameters within SAFT-$\gamma$ Mie which we discuss here:

- For inorganic ions ($Na^+$ and $Cl^-$), no cross-interaction parameters between them and the various mRNA functional groups have been regressed in the literature. This is primarily due to a lack of experimental data, but also because of the low solubility of such species in these conditions. Nonetheless, one can anticipate that these interactions will be extremely unfavorable. As such, for simplicity, we assign all cross-interaction parameters between mRNA groups and inorganic ions to be zero ($\epsilon_{ij} = 0$).
- To extrapolate to other alcohols, it is assumed that cross-interaction parameters of the primary alcohol group is identical to that in secondary alcohols and methanol. It is generally expected that these parameters should be very similar. This assumption is intended to avoid reliance on additional experimental data.
- While a single functional group may exist multiple times in a given species, its chemical environment is expected to have an influence on its cross-interaction parameters. For example, a nitrogen group within a ring is expected to interact differently if it is part of a six-membered ring versus a five-membered ring. We account for these differences through second-order groups where the same self-interaction parameters are used while having different cross-interaction parameters. Groups used in the work have been listed in table SI.1.

- We also note that two groups (cyclic ketones and fused aromatic groups) were fitted by other authors[1, 2] in parallel to this work. In principle, these groups should still be compatible with those presented in this work. However, for the purposes of modeling mRNA solubility, we recommend using only the groups provided in this work.

Table SI.1: Molecular groups (and subgroups) available in the SAFT-$\gamma$ Mie approach used in this work. Additional context for each group, as well as their respective sources, have been provided.

| Main group | Subgroup | Context | Source |
|---|---|---|---|
| H2O | H2O | Water | [3] |
| CH3COCH3 | CH3COCH3 | Acetone | [4] |
| CH3 | CH3 | Terminal carbon group | [5] |
| CH2 | CH2 | Linear carbon group | [5] |
| CH2OE | CH2OE | Linear carbon group in PEG | [6] |
| CH3OH | CH3OH | Methanol | [3] |
| CH2OH | CH2OH | Primary alcohol group bonded to a carbon group | [7] |
| CHOH | CHOH | Secondary alcohol group bonded to a carbon group | [8] |
| aCH | aCH | Aromatic carbon group | [9] |
| | aCH_5 | Aromatic carbon group on a 5-membered ring | This work |
| aCCH2 | aCCH2 | Methylene group bonded to an aromatic carbon group | [4] |
| aCCH3 | aCCH3 | Methyl group bonded to an aromatic carbon group | [4] |
| | aCCH3_5 | Methyl group bonded to an aromatic carbon group on a 5-membered ring | This work |
| afC | afC | Aromatic carbon between two fused rings | This work* |
| | afCp | Aromatic carbon between two fused rings where one is a 5-membered ring | This work |
| aN | aN | Aromatic nitrogen group | This work |
| | aN_5 | Aromatic nitrogen group on a 5-membered ring | This work |
| | aN_2 | Aromatic nitrogen group when two are present on an aromatic ring. (e.g. adenine, pyrimidine) | This work |
| aCNH2 | aCNH2 | Amine bonded to an aromatic carbon | This work |
| cCH2 | cCH2 | Cyclic carbon group | [9] |
| | cCH2_5 | Cyclic carbon group on a 5-membered ring | This work |
| cNH | cNH | Amine group on a ring | [10] |
| | cNH_5 | Amine group on a 5-membered ring | This work |
| cN | cN | Thrice bonded nItrogen group on a cyclic ring | [10] |
| | cN_5 | Thrice bonded nItrogen group on a 5-membered ring | This work |
| cO | cO | PEG ether group | [6] |
| cyO | cyO | Cyclic ether group | This work |
| cCHOH | cCHOH | Hydroxyl bonded to a cyclic carbon | [11] |
| | cCHOHn | Hydroxyl bonded to a cyclic carbon on a nucleotide | This work |
| cC=O | cC=O | Cyclic ketone group | This work† |
| | cC=O_2 | Cyclic ketone group when two are present on an aromatic ring (e.g. guanosine, theophylline, theobromine, caffeine) | This work |
| Na+ | Na+ | Sodium ion | [12] |
| Cl- | Cl- | Chloride ion | [12] |
| NH4+ | NH4+ | Ammonium ion | This work |
| PO4- | PO4- | Phosphate ion (mRNA) | This work |
| | PO43- | Phosphate ion (free) | This work |
| COO- | COO- | Ester ion group | [13] |

*Alternative parameters available at: [1]
†Alternative parameters available at: [2]

### 1 Supplementary Figures

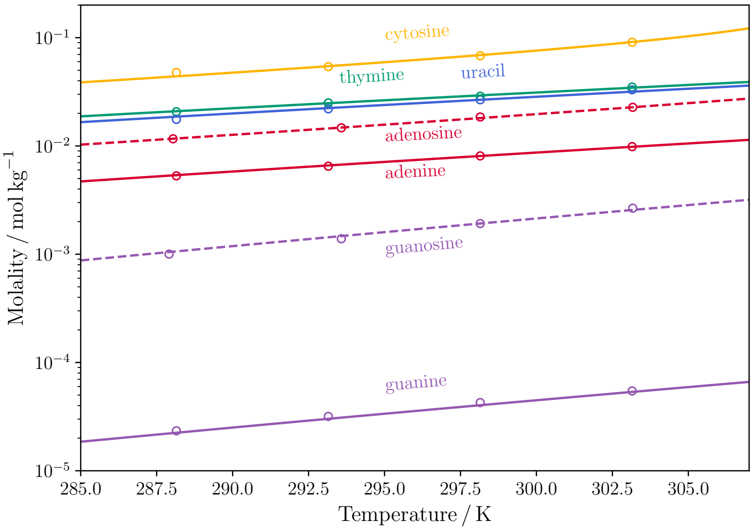


Figure SI.2: Solubility of various nucleobases and nucleosides in water with varying temperature. Lines represent estimates from the SAFT-$\gamma$ Mie equation and symbols represent experimental data.

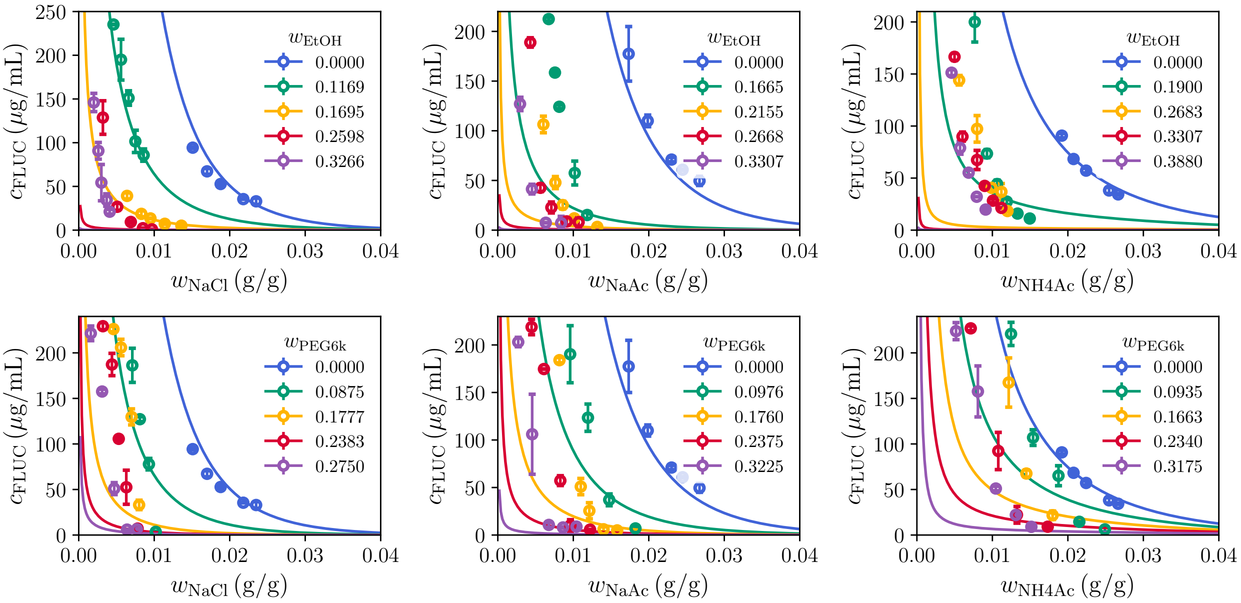


Figure SI.3: Solubility of COVID sequence at 277.15 K with varying salt concentration in both ethanol and PEG6k solvent conditions. Lines represent estimates from the SAFT-$\gamma$ Mie equation and symbols represent experimental data.

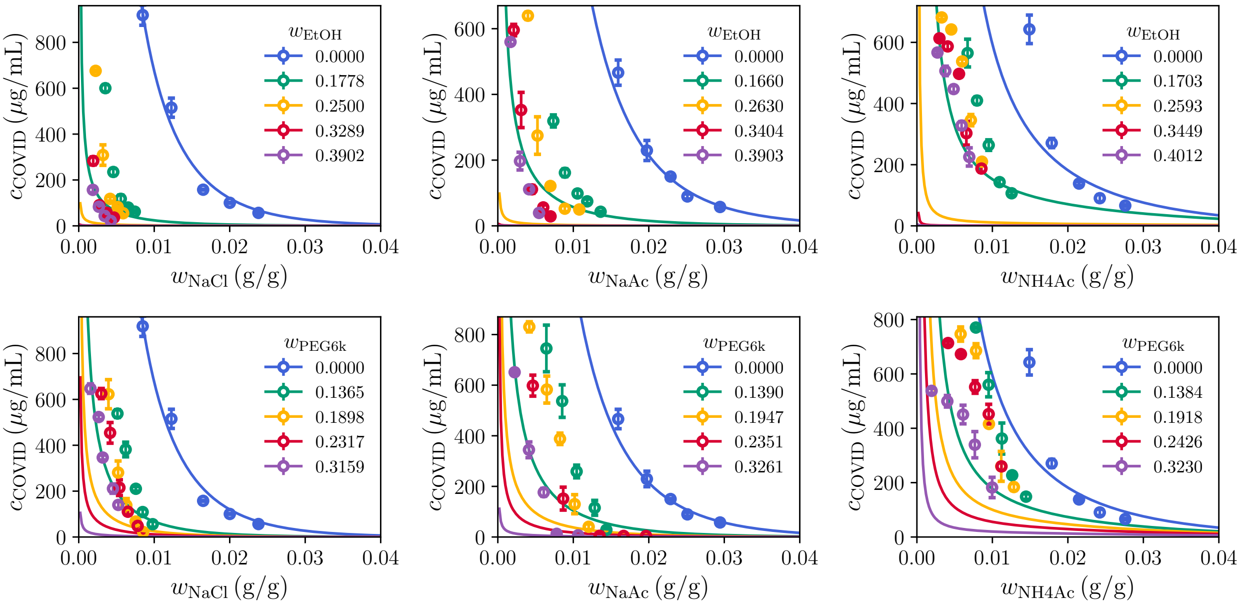


Figure SI.4: Solubility of COVID sequence at 277.15 K with varying salt concentration in both ethanol and PEG6k solvent conditions. Lines represent estimates from the SAFT-$\gamma$ Mie equation and symbols represent experimental data.
